# Microfluidic Rapid and Autonomous Analytical Device (microRAAD) to Detect HIV from Whole Blood Samples

**DOI:** 10.1101/582999

**Authors:** Elizabeth A. Phillips, Taylor J. Moehling, Karin F.K. Ejendal, Orlando S. Hoilett, Kristin M. Byers, Laud Anthony Basing, Lauren A. Jankowski, Jackson B. Bennett, Li-Kai Lin, Lia A. Stanciu, Jacqueline C. Linnes

## Abstract

Early Human Immunodeficiency Virus (HIV) testing is critical to preventing transmission and providing treatment to HIV-positive individuals, yet an estimated 30% of HIV-positive individuals do not know their status because of barriers to early diagnosis. Readily accessible, highly sensitive, and rapid diagnostic tests would enable patients’ prompt treatment with anti-retroviral therapies and reduce transmission. However, existing HIV diagnostic technologies either do not detect early stages of infection or require multiple days of laboratory processing, delaying notification of patients’ status.

Molecular techniques that amplify HIV RNA can detect the earliest stages of infection, within 8-10 days after transmission. However, most of these molecular assays require cold-chain storage of reagents, significant sample preparation, and extensive laboratory infrastructure. To achieve early detection, we developed a reverse transcription loop-mediated isothermal amplification (RT-LAMP) assay with a limit of detection of 10 HIV-1 RNA copies visualized by eye using a lateral flow immunoassay. To demonstrate automated sample-to-answer detection of HIV, we incorporate dried amplification reagents and wax valves in low-cost substrates with resistive heating elements and circuitry. By combining controlled heating with paper’s capillary flow, our assembled device automatically isolates viral particles from human blood samples, amplifies HIV-1 RNA, and transports products to a detection zone. We determine that as few as 10^5^ HIV-1 viral particles can be separated from whole blood, amplified, and visually detected within 90 minutes of sample addition into our Microfluidic Rapid and Autonomous Analysis Device (microRAAD). The low-cost and automated attributes of microRAAD demonstrate its utility as a point-of-care testing platform.

## Introduction

Despite the effectiveness of antiretroviral therapy (ART) to suppress viral loads and decrease HIV-related mortality, HIV remains a global epidemic. The World Health Organization (WHO) estimates that of the 36.7 million people currently living with HIV worldwide, only 48% are being treated (World Health Organization, 2018). Early diagnosis of HIV decreases mortality and morbidity by initiating early patient treatment (Palella et al., 2003). Given that only 70% of the HIV-positive population knows their status, improved HIV diagnosis at the point of care (POC) have the potential to increase distribution of proper treatment and reduce transmission to others (WHO, 2016; World Health Organization, 2018).

HIV screening is currently performed using commercially-available rapid diagnostic tests (RDT), typically based on lateral flow immunoassay (LFIA) technology that detects HIV antibodies from oral fluid or capillary blood. The low sensitivity during the pre-seroconversion phase of the first 4 weeks of infection and frequent false negatives require that RDT results are confirmed by a second or even third laboratory-based assay (Parekh et al., 2019). Fourth and fifth generation combined antibody and HIV p24 antigen tests are more sensitive than antibody-detecting RDTs, however, these must be performed in a laboratory with luminescence-measuring instrumentation because poor signal intensity prohibits visual detection by eye (Alexander, 2016). The multi-day delay of diagnosis due to this laboratory-based testing significantly impairs an HIV-positive patient’s prompt treatment (WHO, 2016).

POC nucleic acid-based diagnostic tests could expedite treatment response for vulnerable and newly infected individuals through early detection of the HIV virus. Reverse transcription polymerase chain reaction (RT-PCR) has been performed in microfluidic-based sample-to-answer devices to amplify HIV RNA spiked into saliva samples. However, the complexity of manufacturing a device to perform sample preparation and cyclical heating often makes it prohibitively expensive for low-resource settings (Chen et al., 2013). While designed for ART failure testing, these sensitive detection systems are not cost-effective for early screening and POC testing because they require expensive supporting sample preparation units, cold-chain storage of reagents, and trained users (Calmy et al., 2007; Parekh et al., 2019). To address these shortcomings, recent efforts have been focused towards developing integrated sample-to-answer nucleic acid analysis devices that can be used by minimally-trained personnel (Choi et al., 2015; Lafleur et al., 2016). There are a few commercial tools for rapid detection of HIV including Cepheid Xpert Qual Assay, Alere q HIV-1/2 Detect, and Diagnostics for the Real World’s Samba II. Although these tests were able to integrate and automate sample preparation, they all require cost-prohibitive (>$17,000 for the instrument and >$17 for the cartridge) benchtop instruments with stable electrical power supply or consumable batteries (Lemaire, 2017).

Recent advances in technologies for point-of-care molecular detection of HIV include several isothermal nucleic acid amplification techniques that could reduce the complexity and therefore cost of a fully integrated testing device (Mauk et al., 2017). One such isothermal amplification method, loop-mediated isothermal amplification (LAMP), provides specific and efficient amplification of target nucleic acids by targeting 8 unique sequences (Nagamine et al., 2002). Operable at a single temperature (most efficiently between 65 and 72 °C) (NEB, n.d.; Rolando et al., 2019), LAMP robustly amplifies even in the presence of complex sample matrices, further reducing sample processing and instrumentation requirements (Clayton et al., 2019; Mori and Notomi, 2009; Phillips et al., 2018). To expedite sample preparation steps, such as reverse transcription (RT) of HIV RNA targets, several groups have demonstrated that RT can be performed using the same assay conditions as LAMP (Curtis et al., 2012; Damhorst et al., 2015; Odari et al., 2015; Rudolph et al., 2015). Gurrala et al. uses RT-LAMP to amplify HIV-1 RNA and produce a pH change that can be measured with their device (Gurrala et al., 2016). The lack of reagent storage and integrated sample preparation, however, decreases the translatability of this and other RT-LAMP devices.

Here we report a fully-integrated sample-to-answer platform that leverages paper membranes’ wicking abilities and size discriminating pores to a) isolate HIV viral particles from human blood samples, b) amplify RNA from the viral particles using pre-dried RT-LAMP reagents that target the highly conserved *gag* gene of HIV-1 and c) automatically transport RT-LAMP amplicons to a downstream LFIA for simple, visual interpretation of results within 90 minutes of sample application. This work demonstrates the potential for early and low-cost HIV detection at the point of care.

## Materials and Methods

### Reagents

Reagents necessary for the RT-LAMP assay included six primers (Integrated DNA Technologies, Skokie, IL), Bst 3.0 polymerase (NEB, Ipswich, MA), deoxynucleotide triphosphates (dNTPs) (Agilent Technologies, Santa Clara, CA), isothermal buffer II (NEB, Ipswich, MA), betaine (Millipore Sigma, Burlington, MA), EvaGreen (VWR International, Radnor, PA), ROX (Thermo Fisher Scientific, Waltham, MA), diethyl pyrocarbonate (DEPC) water (Invitrogen, Carlsbad, CA), and human whole blood collected in sodium citrate (Innovative Research, Novi, MI).

Template used in the experiments below included purified genomic RNA from HIV-1 (ATCC, Manassas, VA), non-infectious HIV-1 virus (AccuSpan, SeraCare Life Sciences, Milford, MA), purified genomic RNA from dengue (DENV) Type 1 (BEI resources, Manassas, VA), and purified RNA from chikungunya (CHIKV) S-27 (BEI resources, Manassas, VA). *Sph*I and *Pst*I restriction enzymes (NEB, Ipswich, MA) and phosphate buffered saline (PBS) (Thermo Fisher Scientific, Waltham, MA) are additional reagents used.

### Reverse Transcription Loop-Mediated Isothermal Amplification (RT-LAMP)

Purified genomic RNA from HIV-1 or non-infectious HIV-1 virus was used as the template in preliminary testing and at specified concentrations in experiments thereafter. LAMP primers were devised using PrimerExplorer v5 to target the *gag* gene of HIV-1, and loop primers were labeled with 6-carboxyfluorescein (FAM) and biotin for detection via commercial LFIA (Ustar Biotechnologies, Hangzhou, China). The primer sequences are provided in Table S1. To allow for both reverse transcription and amplification of the HIV-1 target, we used Bst 3.0 polymerase which includes reverse transcriptase capabilities and the buffers and dyes listed in Table S2 in a 25 µL reaction. 2-4 µL of template or negative control (DEPC water) were added directly to the RT-LAMP master mix prior to heating. A range of temperatures, 58°C – 74°C, were tested to determine the optimal assay temperature. After 25 minutes of heating at various temperatures, RT-LAMP amplicons were added to LFIAs for analysis. Hereafter, RT-LAMP was performed at 65°C for 60 minutes using an Applied Biosystems 7500 Real-Time PCR System (Foster City, CA). The specificity of the target sequence to HIV-1 was verified by BLAST as well as experimentally using 10^5^ copies/reaction of genomic RNA from DENV Type 1 and CHIKV S-27. Further, to confirm the identity of the amplified product, the RT-LAMP amplicons were subjected to restriction enzyme digest with *Sph*I and *Pst*I for 1 hour at 37°C and the digested segments were confirmed via 2% agarose gel electrophoresis.

For limit of detection (LOD) experiments, RNA or virus template was prepared by performing 10-fold serial dilutions in either DEPC water (RNA) or AccuSpan serum (virus). For analysis of RT-LAMP in various biological sample matrices, increasing amounts of human whole blood or plasma was added into the master mix. Real-time fluorescence data of EvaGreen intercalating dye and ROX reference dye was monitored to confirm the amplification progress. RT-LAMP amplicons were characterized via LFIA and confirmed via gel electrophoresis using a 2% agarose gel run at 100 V for 60 minutes, stained with ethidium bromide, and imaged using an ultraviolet light gel imaging system (c400, Azure Biosystems, Dublin, CA).

### RT-LAMP Reagent Drying and Storage

Reagents and conditions for drying and storage were modified and adapted from a published protocol (Hayashida et al., 2015). The RT-LAMP reagents were deposited, dried, and stored at room temperature, eliminating the need for cold-chain storage (−20°C) and improving the portability of the device. The primer mixture (Table S3) containing primers, sucrose, glycerol, and Triton X-100 was deposited by hand on 1 cm wide polyethylene terephthalate (PET) film (Apollo, Lake Zurich, IL) in two parallel lines (Figure S1) at approximately 1.83 µL/cm. After drying in a sterile biosafety cabinet under continuous air flow for 60 minutes at room temperature, the enzyme mixture containing Bst 3.0 polymerase, sucrose, and dNTPs was deposited directly on top of the dried primers in parallel lines at approximately 1.19 µL/cm and set out to dry for another 60 minutes. The PET with dried reagents was cut into 1 × 1 cm pieces, corresponding to one 25 µL reaction.

The reagents for the stability studies were packaged after deposition and initial drying and stored in opaque Mylar bags with silica gel desiccant (Uline, Pleasant Prairie, WI) at room temperature for 3 weeks. The dried RT-LAMP reagents were rehydrated with buffer (Table S3) and virus (positive samples) or DEPC water (negative controls) in tubes or in the polyether sulfone (PES) amplification zone within the integrated device.

### Blood Separation and Virus Capture in Paper Membranes

As an initial proof of concept of size-based separation, two sizes of fluorescent nanoparticles (Bangs Laboratories, Fishers, IN) were used in vertical flow filtration to allow quantification of the membrane capture efficiency. The 0.11 µm diameter Dragon Green (Ex480/Em520 nm) nanoparticles represented HIV-1 virus and the 7.32 µm diameter Suncoast Yellow (Ex540/Em600 nm) nanoparticles represented red blood cells. The fluorescent nanoparticles were diluted in PBS according to manufacturer’s instructions. A calibration curve correlating nanoparticle concentration to fluorescence was created by serially diluting both nanoparticle solutions and measuring dilutions in a SpectraMax M5 microplate reader (Molecular Devices LLC, San Jose, CA) at excitation of 480 nm or 540 nm for the 0.11 µm and 7.32 µm nanoparticles, respectively. A 7 mm hole punch was used to cut pieces of blood separation membrane (MF1, GE Healthcare, Chicago, IL) and amplification membrane (0.22 µm PES, Millipore Sigma, Burlington, MA). The cut membranes were sandwiched between two O-rings and placed into a commercial miniprep spin column (Figure S2) (Qiagen, Hilden, Germany). The spin column was then placed into a clear 2 mL collection tube.

One hundred fifty (150) µL of either the 0.11 µm or 7.32 µm nanoparticles was pipetted into the spin column containing the membrane of interest. The tubes were then centrifuged for 60 seconds at 0.5 rcf and the fluorescence of the eluent was measured and compared to the calibration curve intensities. Fluorescence of unfiltered nanoparticles was measured and used as the baseline to calculate the proportion of nanoparticles that passed through the membrane.

MF1 and 0.22 µm PES membranes were then used to show size-based capture in a lateral flow format. The 0.22 µm PES (2.5 cm × 1 cm) was overlapped with the MF1 membrane (1 cm × 1 cm) to form the amplification and filtering segments of the integrated device (Figure S10). First, a 100 µL solution containing approximately 7 × 10^5^ of 0.11 µm nanoparticles, 230 of 7.32 µm nanoparticles, and deionized water were mixed to form the nanoparticle mixture. Thirty (30) µL of the nanoparticle solution was pipetted onto the assembled MF1/PES membranes, followed by a 30 µL PBS wash. The nanoparticles in the membranes were immediately imaged at 40X magnification with an inverted Axio Observer Z1 Fluorescent microscope and ZenPro software (Carl Zeiss Microscopy, Thornwood, NY) and using a Rhodamine dye filter cube for the 7.32 µm nanoparticles and an Alexa Fluor 488 dye filter cube for the 0.11 µm nanoparticles.

To demonstrate nucleic acid amplification of viral particles after separation from blood cells in the lateral flow format, MF1 and 0.22 µm PES membranes were overlapped as in the nanoparticle lateral flow. 1.2 µL of HIV-1 was mixed with 12 µL of human whole blood (amounting to 2.3 × 10^7^ virus copies/mL) and deposited onto the MF1 membrane of the MF1/PES assembly, followed by a 61.8 µL wash of rehydrating mixture (Table S3). After 1 minute of capillary flow, the PES was removed from the assembly and added into a PCR tube with 23 µL of the enzyme and primer mixtures. The samples and a positive amplification control (reaction without blood or membrane) were amplified for 60 minutes at 65°C. Amplification was confirmed by placing the PES membranes and control reaction into wells of an agarose gel and performing gel electrophoresis. The remaining solution in the PCR tube that had not saturated the PES membrane was added to a LFIA with 40 µL of wash buffer.

### Resistive Heating

Resistive heating elements were fabricated by printing JS-B40G Nanosilver Ink (Novacentrix, Austin, TX) onto a Kapton substrate (Kapton HN Semi-Clear Amber Film, 5 mm × 125 μm) using a Dimatix Materials Printer DMP-2850 (Fujifilm Dimatix Inc., Santa Clara, CA) with a drop spacing of 35 µm and a platen temperature of 50°C. Optimization and characterization were performed previously. The printed traces were then cured in an oven at 400°C for 10 minutes. Individual traces were measured post-curing with a handheld multimeter to determine the average electrical resistance of the heating element. The design in Figure S3 provided even heating of the amplification zone with an average resistance of 5 Ohms and required an average of 240 mW to reach 65°C, the temperature necessary for the RT-LAMP assay. The heating elements used to actuate the wax valves also had an average resistance of 5 Ohms and heated to 80°C using an average of 440 mW. The heat produced by the silver trace dissipates through the Kapton substrate and into the membranes above. The insulation required to maintain steady heating is provided by an acrylic lid and plastic housing (Figure 1A).

**Figure 1.**
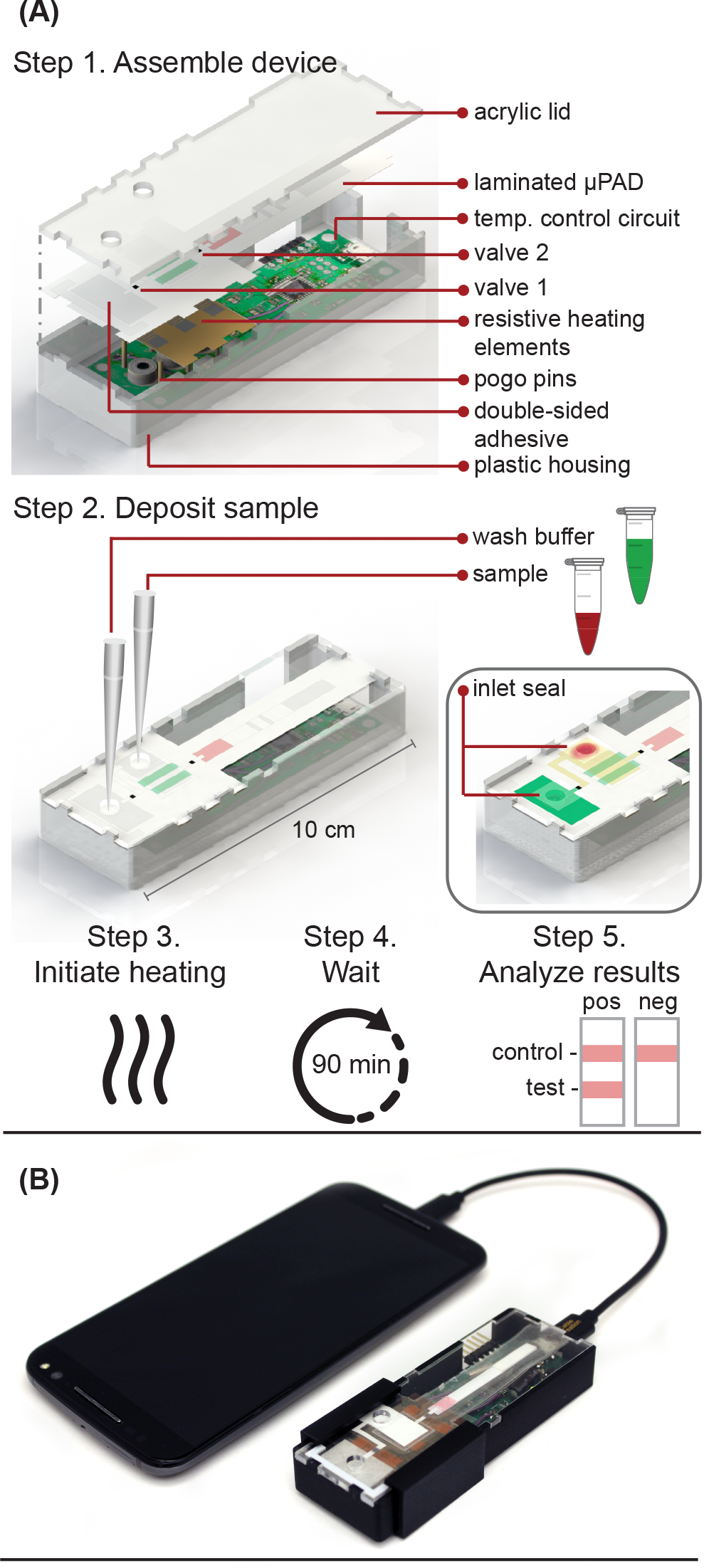
**(A)** Schematic of microRAAD for HIV testing work flow in which user 1) assembles consumable μPAD into plastic housing with reusable resistive heating elements and temperature control circuit, 2) deposits sample and wash buffer into inlets and seals with tape to minimize evaporation, 3) initiates heating by connecting to phone, 4) waits 90 minutes for automated fluid delivery and sample incubation in μPAD, and 5) analyzes results of lateral flow immunoassay. **(B)** Photo of microRAAD connected to phone to power the temperature control circuit.

### Temperature Control Circuit

In the integrated microRAAD, temperature regulation of the resistive heating elements was achieved using a miniaturized, custom-designed electronic device. It was constructed to individually monitor and regulate the temperature of each of three heating zones: one heating zone was dedicated to amplification and two heating zones were dedicated to actuating the wax valves. The temperature control circuit was equipped with a microcontroller (ATMEGA328P-AUR, Microchip Technology, Chandler, AZ), three non-contact infrared (IR) temperature sensors (MLX90614ESF-BAA-000-TU, Melexis Technologies NV, Ypres, Belgium), and three transistor-based current drivers (IRLML6244TRPBF, Infineon Technologies, Neubiberg, Germany) driven by three separate 10-bit digital-to-analog converters (DAC6311, Texas Instruments, Dallas, TX) for precise monitoring and control of current delivered to the resistive heating elements. Six pogo pins (0907-1-15-20-75-14-11-0, Mill-Max Manufacturing Corporation, Oyster Bay, NY), two for each resistive heating element, were used to complete the circuit between the current drivers and the resistive heating elements placed between the circuit board and the microfluidic paper analytical device (µPAD, Figure 1A). The microcontroller was loaded with an Arduino bootloader (Arduino Pro Mini, 3.3V, 8MHz, SparkFun Electronics, Boulder, CO) and programmed using the Arduino development environment (Arduino IDE v.1.8.8) over a Universal Serial Bus (USB) connection through a Future Technology Devices International (FTDI) serial-to-USB interface (FTDI Serial TTL-232 USB Cable, Adafruit Industries, New York City, NY). All electronic components were sourced from Digi-Key Electronics (Thief River Falls, MN). The circuit boards were fabricated by PCBWay (Shenzhen, China) and were assembled in-house using standard soldering equipment.

The microcontroller was programmed to implement a proportional integral differential (PID) algorithm for maintaining temperature of the amplification zones within the user-specified set points (65°C for the amplification and 80°C for each wax valve). PID algorithms are popular closed-loop feedback mechanisms due to their simplicity and effectiveness, making them the de facto choice for portable, low computing power electronic devices (A□stro□m and Murray, 2008; Ziegler and Nichols, 1993). The microcontroller sampled the temperature of each zone (sampling at 16.2 Hz for the amplification zone and 13.5 Hz for the valves) using the non-contact IR sensors and compared the measured temperature to the user-specified set point in order to determine the error value, *e(t)* (the difference between the measured value and the set point as depicted in Equation (1)). The device then computes the proportional, integral, and derivative terms of the PID algorithm. The proportional term represents the difference between the set point and the measured value, multiplied by the proportional gain, *K*_*p*_ = 0.05. The integral term is the cumulative error and is computed by summing the integral of the current error with the previous errors, multiplied by the integral gain, *K*_*i*_ = 0.0001. The derivative term represents the change in the error since the last measurement, multiplied by the derivative gain, *K*_*d*_ = 0.1.

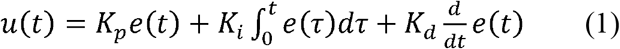

The microcontroller then adjusts the current delivered to the resistive heating elements accordingly. The algorithm is designed to achieve accurate temperature regulation within 0.7°C of the set point, while minimizing overshoot (1°C). A serial Arduino interface was enabled between the circuit and a computer allowing real-time monitoring of experimental parameters, including temperature of each zone, time elapsed, and power consumption. We verified the temperature with both an infrared camera (FLIR Systems, Wilsonville, OR) and K-type thermocouple measurement of the top and bottom surfaces of the μPAD using a portable temperature data logger (RDXL4SD, OMEGA Engineering, Norwalk, CT).

In the final implementation, the temperature control circuit was powered through a USB port using a USB On-The-Go (USB OTG) enabled cellphone (Samsung Galaxy J3 Luna, Android Version 7.0), which removed the need for the computer, increased portability, and ensured fully automated control of the integrated device without the need for user intervention.

### Integrated microRAAD

The microRAAD for HIV detection is composed of the reusable temperature control circuit and silver ink resistive heating elements and a single-use, laminated μPAD, all contained in a plastic housing (Figure 1A). The base of the plastic housing was designed in SolidWorks and 3D printed on a Fortus 380c printer (Stratasys, Eden Prairie, MN). The 0.08” thick acrylic lid of the housing (model #11G0670A, Shape Products Menomonie, WI) and components of the single-use μPAD (Table S4) were designed in Adobe Illustrator and cut with a CO_2_ laser (VLS 3.5, Universal Laser Systems, Scottsdale, AZ). Valve strips were prepared by printing 1.25 mm wide lines of solid wax-ink (Black ColorQube ink, Xerox, Norwalk, CT) onto cellulose membranes (Chr1, GE Healthcare, Pittsburgh, PA) using a Xerox ColorQube 8570 (Norwalk, CT). Membranes were then heated for twelve minutes at 85°C in a table-top oven (VWR, Radnor, PA) and cut into strips with the laser cutter to create closed valve strips (Phillips et al., 2016). Commercially available LFIAs were modified for microRAAD by cutting off the sample pad. PET squares were sterilized with 70% ethanol. Once prepared, all components were hand assembled (Table S4 and Figure S4) and laminated with pressure sensitive Self-Seal adhesive (GBC, Lake Zurich, IL) to minimize evaporation during the assay. Seventy-five (75) μL of prepared RT-LAMP master mix or rehydrating mixture (Table S3) containing HIV-1 virus (at a final concentration of 4 × 10^6^ virus copies/mL) were loaded into the sample inlet of the μPAD and sealed with a 1×1 cm square of Self-Seal to minimize evaporation and contamination. When testing whole blood samples spiked with HIV-1 virus, 1.2 µL of HIV-1 was mixed with 12 µL of human whole blood (amounting to 2.3 × 10^7^ virus copies/mL) and loaded into the sample inlet, followed by a 61.8 µL addition of rehydrating mixture. The loaded μPAD was then adhered to the acrylic lid with double-sided adhesive at the wash inlet. Resistive heating elements were adhered to the backside of the μPAD, aligned with the two valves and amplification zone, and faced such that the silver traces would contact the pogo pins of the temperature control circuit inside the plastic housing. Two plastic brackets were slid over the acrylic lid and plastic housing to ensure proper contact within microRAAD. Finally, 130 μL of green food coloring solution (for visualization of flow) were added to the wash inlet and sealed. Heating was initiated via the serial interface between the computer and the temperature control circuit: 1) 65 °C for the middle resistive heating element (amplification) for 60 minutes and 2) 80 °C for the outer resistive heating elements (valves) for 2 minutes. After 30 minutes of development (1.5 hours after initiating the heating), the LFIA was imaged using a desktop scanner for analysis (Epson, Suwa, Japan).

### LFIA Quantification and Statistical Analysis

All RT-LAMP amplicons were characterized via LFIA. The LFIAs were scanned at least 30 minutes after initial sample addition using an Epson V850 Pro Scanner. The test band was quantified using a custom MATLAB script that averages the grey-scale pixel intensity of the test band and subtracts out the average background pixel intensity 25 pixels below the test band (Holstein, 2015). To determine the limit of detection, a one-way ANOVA post-hoc Dunnett’s was performed with multiple comparisons against the negative control with no template with a 95% confidence interval. A Student’s unpaired, two-sided t-test with a 95% confidence interval was used when comparing the negative control and positive samples during the initial testing of 21-day dried reagents and to determine significance between control and positive samples detected in the integrated device.

## Results and Discussion

### RT-LAMP Optimization

We designed new primers to specifically target the 201bp sequence in the *gag* gene after experiencing slow amplification and poor signal generation from several published HIV LAMP primer sets. There is one copy of the *gag* gene in the HIV-1 genome and two copies of the genome per viral particle (Sundquist and Kräusslich, 2012). Optimized assay conditions are shown in Table S2 and provided the greatest amplification time difference between positive and negative samples. The optimal RT-LAMP assay temperature for this primer set is 65°C as this temperature yielded the strongest test band intensity on LFIAs illustrated in Figure S5. To assess specificity of the RT-LAMP assay, primers were tested with DENV and CHIKV RNA. As seen in Figure S6, there was no cross-reactivity of the HIV LAMP primers with these other RNA-based viral pathogens. Further confirmation of specificity was provided by performing enzymatic digestions of the amplified product with either *Sph*I or *Pst*I. As predicted by LAMP restriction digest fragment analysis (Reisle, 2015), digestion with either of these enzymes resulted in smaller fragments compared to the undigested product, with the *Pst*I digested product collapsing to the shortest fragment seen on the agarose gel in Figure S7. Taken together, these experiments suggest that the designed primer set targets the intended region in the *gag* gene and that this target amplification is specific to HIV-1.

### RT-LAMP Characterization

RT-LAMP was first performed with 2 µL HIV-1 RNA at concentrations ranging from 10^1^ – 10^6^ RNA copies/reaction (n=3). As expected, the real time fluorescence data displayed faster amplification for samples with higher initial concentrations of RNA (data not shown). Agarose gel electrophoresis confirmed amplification of the RNA after the 60-minute assay (Figure 2A). The RT-LAMP amplicons were also added to LFIAs and the test band intensity was quantified to determine the LOD. Statistically significant differences between the test band intensity of the negative control (0) compared to 10^2^ - 10^6^ RNA copies/reaction (p-value < 0.01) (Figure 2A) demonstrate an assay LOD of 100 copies of HIV-1 RNA/reaction. Other groups have reported a comparable LOD, ranging from 30 – 250 RNA copies/reaction for their HIV RT-LAMP assay (Curtis et al., 2018; Odari et al., 2015; Rudolph et al., 2015).

**Figure 2.**
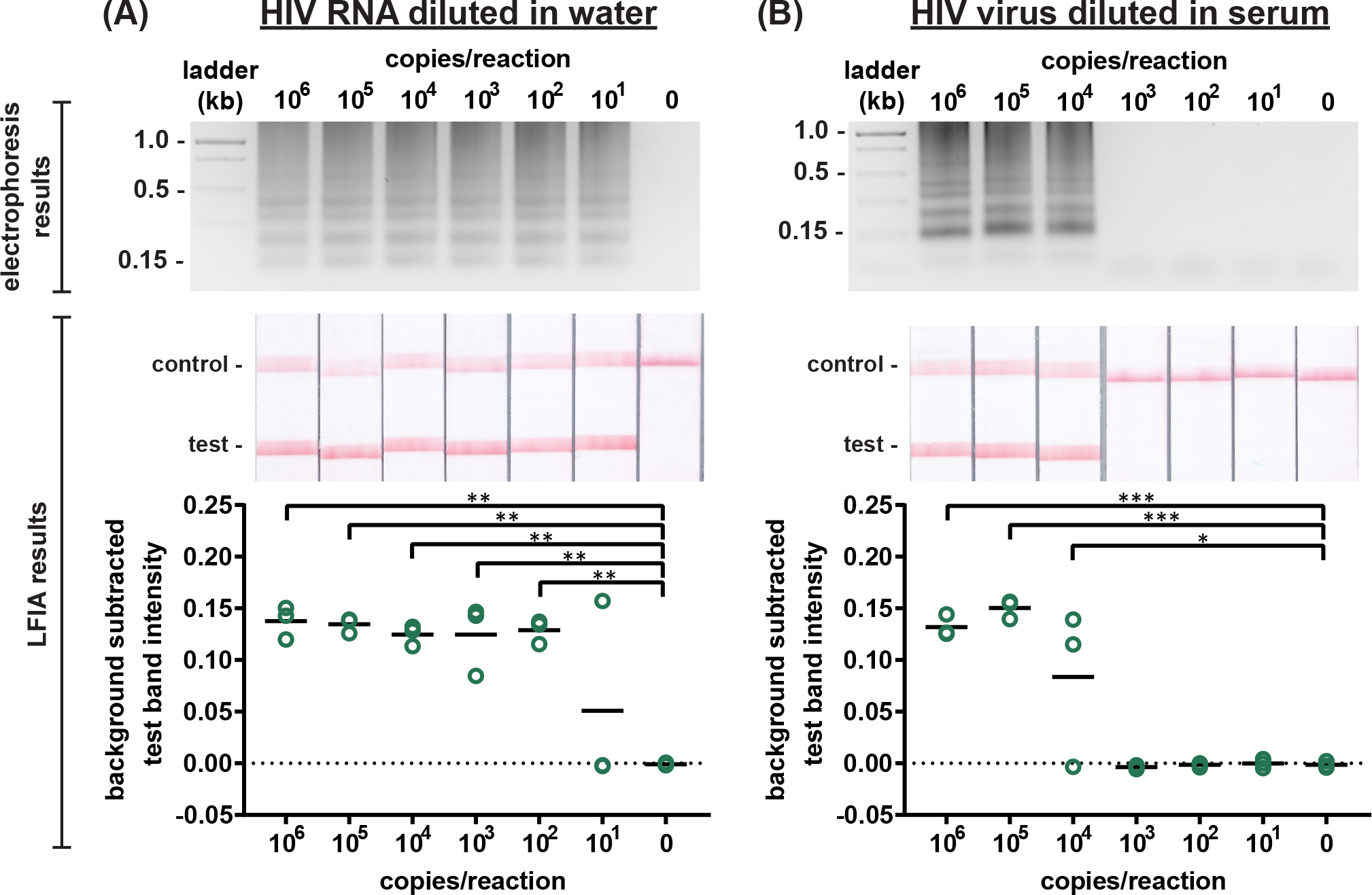
Detection of HIV RNA and virus amplified by RT-LAMP. Electrophoresis gels verifying amplification (top, contrast increased for visualization), LFIA test results (middle), and LFIA test band quantification (bottom). **(A)** Labeled RT-LAMP amplification products are visually detectable from as few as 10 copies of HIV RNA diluted in water. **(B)** Labeled RT-LAMP amplification products are visually detectable from as few as 10,000 HIV viral particles when reactions contain 16% serum. n=3, replicates indicated by each circle; *** indicates p ≤ 0.001; ** indicates p ≤ 0.01; * indicates p ≤ 0.05.

Next, we conducted the RT-LAMP assay on whole HIV-1 virus to ensure sufficient viral lysis at the 65°C assay temperature. Viral lysis is necessary to release the RNA for amplification. Since the osmotic pressure gradient would prematurely lyse the virus, amplification of virus diluted in water was not tested, as was done with the RNA. Instead, we performed RT-LAMP of HIV-1 virus spiked into reactions containing 16% serum at concentrations of 10^1^ – 10^6^ virus copies/reaction (n=3). LFIA analysis showed a statistically significant difference between the test band intensity of 10^4^, 10^5^, and 10^6^ virus copies/reaction compared to the negative control (0) (p-value < 0.05 and p-value < 0.001, respectively) (Figure 2B) and the agarose gel supported this conclusion. Therefore, the limit of detection of the assay with HIV-1 virus in 16% serum is 10^4^ virus copies/reaction. While this LOD is 100X higher than purified RNA, this is, to our knowledge, the first demonstration of an HIV RT-LAMP assay without separate RNA extraction and purification, vastly simplifying the sample preparation required for detection of viruses in complex matrices.

Next, we evaluated the robustness of the RT-LAMP assay in the more complex matrices of human plasma and human whole blood. The RT-LAMP assay performed in 16% plasma demonstrated similar results as the experiments in 16% serum (data not shown). Further, HIV-1 virus spiked into RT-LAMP reactions at a concentration of 10^5^ virus copies/reaction with increasing percentages of whole blood (0-30%) was detectable via gel electrophoresis in up to 15% whole blood while LFIA detected HIV-1 in up to 20% whole blood (Figure S8). In agreement with our observations, other groups have also noticed that LFIA visualization can be more sensitive than gel electrophoresis (Zhang et al., 2017). In the integrated microRAAD device, MF1 will capture the red and white blood cells, allowing the virus to flow into the amplification zone (Figure 1A). Therefore, while whole blood is not intended to be in the amplification zone, our results indicate an assay tolerance up to 20% in case of poor MF1 capture efficiency or blood cell hemolysis.

### RT-LAMP with Dried Reagents

Prior to experimentation, we wanted to determine if the reagent drying setup utilizing parallel lines and layering permits adequate diffusion of the reconstituted reagents into the PES amplification zone using the Stokes-Einstein equation (Einstein, 1905) and Renkin equation (Fournier, 2012; Renkin, 1954). The hydrodynamic radius, *α*, was calculated for the largest molecule in each the primer and enzyme mixture, which was the FIP primer and Bst 3.0 polymerase, respectively. Using Equation (2), we then calculated the bulk diffusivity, *D*, for each molecule where R is the gas constant, T is temperature, *η* is viscosity, and N_A_ is Avogadro’s number.

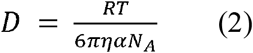

Since the diffusion occurs in a porous membrane, we then calculated pore diffusivity, *D*_*m*_, using Equation (3).

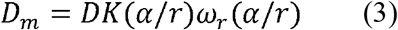

Where *r* is the pore radius, *K(a/r)*, the partition coefficient, is equal to (1 - *α*/*r*)^2^, and the hydrodynamic drag, *ω*_*r*_(*α*/*r*), is equal to [1 - 2.1(*α*/*r*) + 2.09(*α*/*r*)^3^ - 0.95(*α*/*r*)^5^]. From the pore diffusivity, the time for reagents to diffuse 1 mm in the porous PES membrane was estimated at 10 minutes or less, which is sufficient for proper mixing of reagents and sample and subsequent amplification.

To ensure robust detection, 21-day dried RT-LAMP reagents were reconstituted with rehydrating mixture and HIV-1 at a concentration of 10^5^ virus copies/reaction in DEPC water. Positive and negative control reactions using freshly prepared reagents were heated simultaneously. After the 60-minute amplification, samples and controls were analyzed via LFIA and gel electrophoresis (n=6, Figure 3A). The LFIA test band intensity of virus samples using 21-day dried reagents was not statistically significantly different than that of the test band of the freshly prepared positive controls, indicating that drying reagents did not damage enzymatic or primer activity. As expected, LFIA results of positive samples were statistically differentiable from the negative samples for both the dried and fresh reagent groups (p-value < 0.001) (Figure 3A).

**Figure 3.**
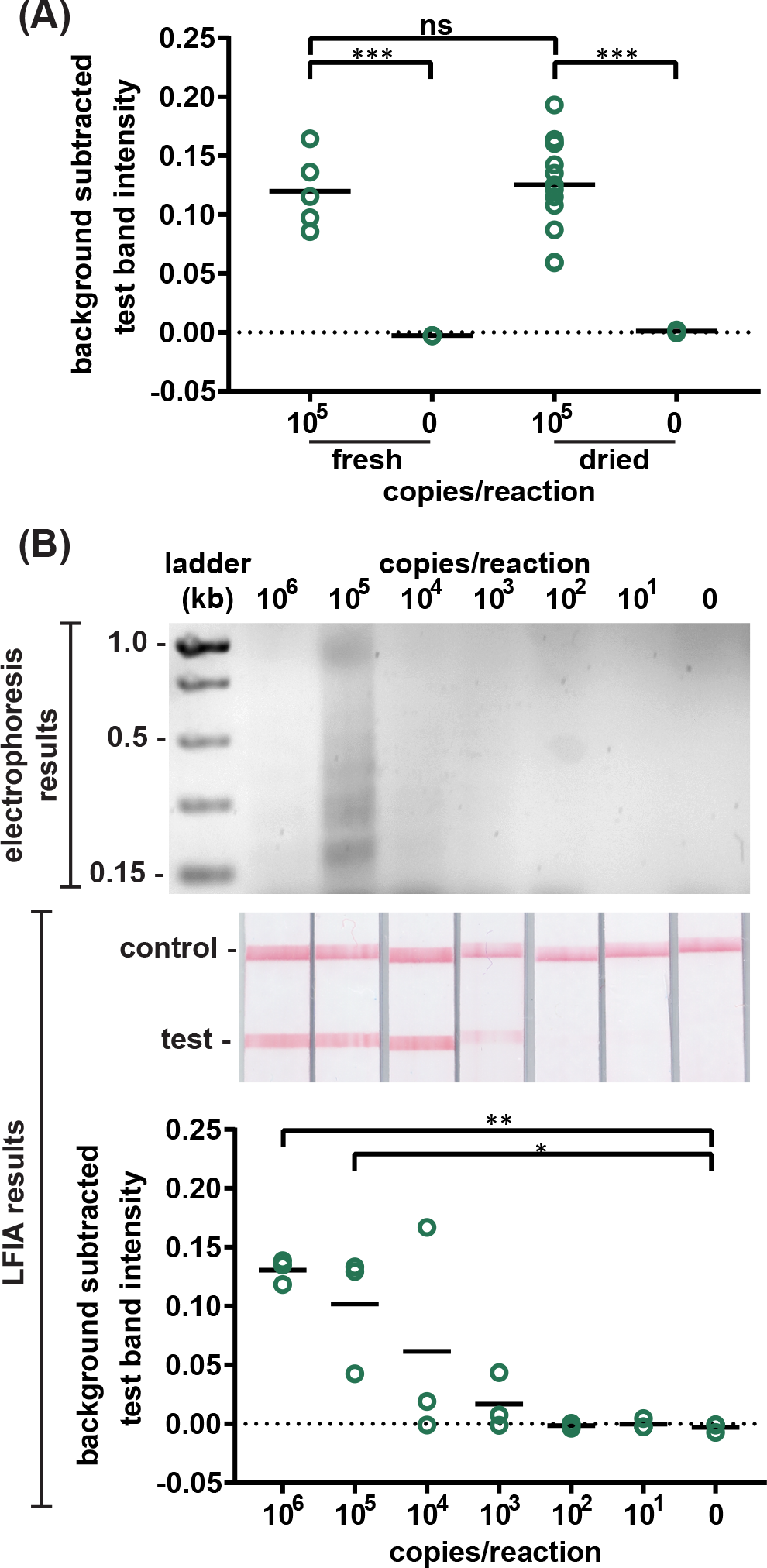
Detection of HIV virus amplified by dried RT-LAMP reagents. **(A)** There is no significant difference in test band intensity of labeled amplification products detected on LFIAs after amplification with fresh RT-LAMP reagents compared to amplification with reagents dried for 21 days. n=5 (fresh), n=13 (dried), circles indicate replicates; *** indicates p ≤ 0.001 **(B)** Labeled RT-LAMP amplification products are visually detectable from as few as 1,000 HIV virus particles when reactions contain 16% serum. Electrophoresis gels verifying amplification (top, contrast increased for visualization), LFIA test results (middle), and LFIA test band quantification (bottom). n=3, circles indicate replicates; ** indicates p ≤ 0.01; * indicates p ≤ 0.05.

To compare the amplification efficiency of dried reagents and freshly prepared reagents, the LOD of HIV-1 in 16% serum was determined using 21-day dried RT-LAMP reagents (n=3). There was a statistically significant difference between the test band intensity of the 10^5^ and 10^6^ virus copies/reaction compared to the negative control (0) (p-value < 0.05 and p-value < 0.01, respectively) (Figure 3B). While not significant, 10^4^ and 10^3^ copies of virus did amplify in some cases; two of three times for 10^4^ and one of three times for 10^3^ virus copies/reaction. There is a slight loss in sensitivity when using the dried reagents (LOD of 10^5^ versus 10^4^ virus copies/reaction), however, the LOD can likely be improved with further assay optimization, such as enzyme selection and primer design. When measuring the LOD with 21-day dried reagents in 16% plasma, we saw a similar outcome compared to 16% serum (data not shown). Another study reported a loss of reaction efficiency compared to freshly prepared controls when using lyophilized HIV RT-LAMP reagents stored at ambient temperature for several hours (Damhorst et al., 2015). However, Hayashida and colleagues established that dried, but not lyophilized, LAMP reagents designed for DNA targets have the same sensitivity as freshly prepared reagents after 7 months of storage at room temperature (Hayashida et al., 2015). These findings in combination with our preliminary studies, shown in Figure S9, give us reason to believe that we can increase the storage time of the HIV RT-LAMP reagents at room temperature beyond 21 days.

### Quantification of Size-Based Capture in Paper Membranes

We developed a simple quantitative method to experimentally test the size-based capture of blood cells and virus in membranes. As seen in Table 1, the 7.32 µm nanoparticles, representative of blood cells, were captured in the MF1 membrane at an efficiency of 98.6% while 30% of the 0.11 µm nanoparticles, representative of the virus, were captured by MF1 (n=3). This implies that MF1 can be used for size-based separation of blood cells from the virus, although some virus will remain in the blood filter thereby reducing detection sensitivity.

**Table 1.** Efficiency of membrane capture of fluorescent nanoparticles. n=3

| <u>Nanoparticle Size</u> | <u>Membrane</u> | <u>Fluorescence (RFU)</u> | <u>Approximate Concentration (nanoparticles/mL)</u> | <u>Capture Efficiency</u> |
| --- | --- | --- | --- | --- |
| **0.11 $\mu\text{m}$** | None | $4007.6 \pm 165.0$ | $4777.1 \pm 1100.7$ | **100%** |
| | MF1 | $2810.9 \pm 193.0$ | $3532.5 \pm 2470.0$ | **30.0%** |
| | 0.22 $\mu\text{m}$ PES | $1986.8 \pm 103.2$ | $2084.1 \pm 169.5$ | **47.6%** |
| **7.32 $\mu\text{m}$** | None | $275.1 \pm 12.2$ | $383.0 \pm 19.3$ | **100%** |
| | MF1 | $2.3 \pm 1.3$ | $4.8 \pm 2.0$ | **98.6%** |
| | 0.22 $\mu\text{m}$ PES | $0.9 \pm 0.2$ | $5.6 \pm 1.1$ | **81.9%** |

Further, 47.6% of the 0.11 µm nanoparticles were trapped in the PES membrane (Table 1, n=3), indicating that the PES captures nearly half of the smaller nanoparticles. Despite the PES membrane having a reported 0.22 µm pore diameter, we suspect that a fraction of smaller diameter nanoparticles were trapped in the PES and the MF1 membranes due to a combination of properties. Because of the membrane heterogeneity, many pores may be smaller than the nominal pore size, allowing nanoparticles to be physically trapped. Furthermore, the tortuosity of the membranes may prevent particle migration through the membrane (Clennell, 1997). Lastly, the proprietary surface chemistries of both membranes may create a slight charge-based attraction that causes nanoparticles to adhere to the membranes. Our experimental results indicate that we can leverage these factors to separate and trap the virus for localized amplification in PES.

### Blood Separation and Virus Capture in Paper Membranes

After quantifying the nanoparticle separation and capture in columns, it was important to demonstrate successful separation and capture in a lateral format. After adding the 7.32 µm and 0.11 µm nanoparticle mixture to the MF1 of the MF1/PES membrane assembly, depicted in Figure S10, separation and capture was qualitatively assessed via fluorescence microscopy. Figure S11 illustrates significant capture of the 7.32 µm nanoparticles in the MF1 membrane at the site where the nanoparticle mixture was added (n=3). Although, some of the 0.11 µm nanoparticles were trapped in the MF1 membrane, the majority were dispersed throughout the PES membrane (Figure S11, n=3), aligning with the vertical flow experiment results and indicating that the virus can be separated from the blood cells and localized to the PES amplification zone.

These proof-of-concept nanoparticle experiments indicated that size-based separation and capture in membranes is possible. The blood cells should be trapped in the MF1 membrane while the virus flows and settles into the PES membrane for subsequent amplification. To test this theory, we spiked HIV-1 virus into human whole blood and added the mixture onto the MF1 membrane which overlapped with the PES membrane and chased the sample with rehydrating mixture. After removing the PES from the MF1/PES assembly and amplifying the trapped virus in the PES membrane, the amplicons were analyzed via LFIA. As depicted in Figure S12, the test band intensity is strong, implying that the virus is dispersed throughout the PES (n=3) yet accessible for amplification. When HIV-1 virus diluted in blood was mixed with the rehydrating buffer and added to the assembly simultaneously, amplification was inconsistent and sometimes completely inhibited because more red blood cells seemingly migrated to the PES (Figure S12). Our successful membrane amplification results are consistent with previous findings that have also shown that LAMP and other isothermal amplification methods can be performed within the PES membrane (Linnes et al., 2016). MF1 membranes inhibited the amplification assay and products extracted from the MF1 were not visible in either the agarose gel or LFIA (data not shown).

### Integration of microRAAD

We verified flow in the assembled μPADs with dyed solutions deposited into the sample and wash inlets and subjected the amplification zone and valves to heating. This initial testing indicated 130 μL and 75 μL were required for the wash and sample, respectively. With too little volume, there was an insufficient pressure gradient to drive flow past opened valves, while with too much volume, the fluid would prematurely leak past the valves. The wash solution is necessary to sufficiently drive the RT-LAMP amplicons from the PES amplification zone into the LFIA for detection. By directing the wash through the MF1, rather than to the PES directly, we ensured that amplicons in the PES migrated to the LFIA before the wash. Moreover, we verified that no capillary leaks occurred between the membranes and lamination by confirming the flow rates of sealed and unsealed membranes were equivalent (data not shown).

For our preliminary studies of HIV detection in microRAAD, we loaded freshly prepared RT-LAMP reagents containing HIV-1 virus onto μPADs and then sealed the sample inlet prior to assembling the μPAD into the plastic housing. This method was used to minimize contamination between experiments, however, we also verified that the sample can be added to the sample inlet once the μPAD is assembled into the plastic housing, which is how we anticipate the device would be used at the point of care. The μPAD containing the sample was then assembled into the plastic housing with the temperature control circuit, loaded with wash buffer, and subjected to localized heating of the amplification zone and valves. We observed the amplification zone reaching 65°C within seconds of initiation and remaining at 65 ± 2°C throughout the 60-minute heating period. The temperature control circuit automatically terminated the amplification zone heating and initiated simultaneous heating of the wax valves. We have previously reported that only 41°C is required to open wax-ink valves prepared in chromatography paper (Phillips et al., 2016), although here we subjected the valves to 80°C to accelerate their opening. Upon initiation of valve heating, the green wash buffer flowed past valve 1 to the MF1 and the RT-LAMP solution flowed past valve 2 into the LFIA portion of the μPAD. Within 5-10 minutes after valves opened, test and control bands were consistently observed on the LFIAs and quantified in Figure S13. In addition to a computer, we found that a fully charged cellphone provided enough current to power the temperature control circuit for the duration of the assay and yielded comparable results (Figure 1B).

Upon verifying the functionality of microRAAD with fresh reagents, we repeated the studies using 21-day dried RT-LAMP reagents and rehydrating buffer containing HIV-1 virus. Samples containing as few as 10^5^ virus copies/reaction resulted in unequivocally positive test bands and samples containing no template (0) yielded negative test results (p-value < 0.05) (Figure 4B). The test band intensity at a concentration of 10^5^ virus copies/reaction using 21-day dried reagents in microRAAD was comparable to the test band intensity of the same concentration in a tube reaction with 21-day dried reagents (Figure 3A).

**Figure 4.**
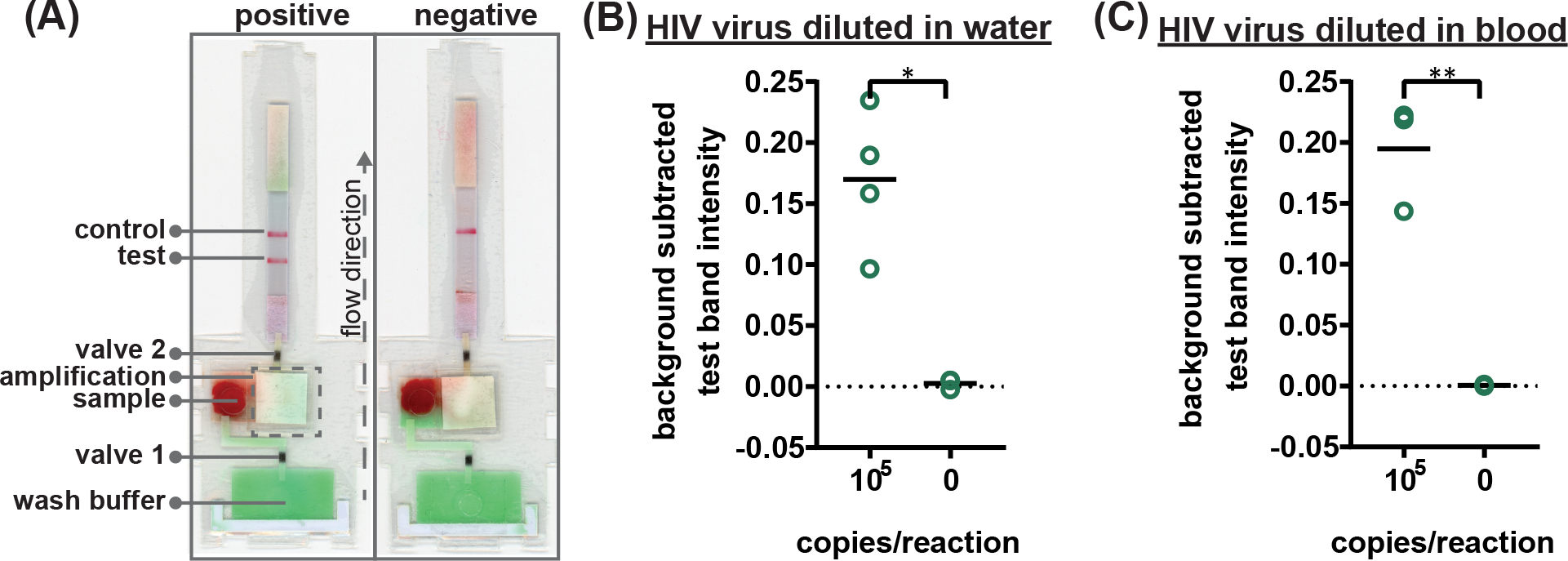
Detection of HIV virus diluted in whole blood on microRAAD with reagents dried for 21 days. **(A)** Representative μPADs imaged 90 minutes after blood (with and without HIV virus) deposited into microRAAD’s sample inlets. After capillary migration of HIV from sample inlet to amplification zone the valves are automatically heated, releasing solution to LFIA for detection. As few as 105 HIV virus copies in **(B)** rehydrating mix alone or **(C)** rehydrating mix with 12 μL of blood are detectable by microRAAD prepared with RT-LAMP reagents dried for 21 days. n=4 (B) and n=3 (C), replicates indicated by each circle; ** indicates p ≤ 0.05 and ** indicates p ≤ 0.01.

Finally, we performed the HIV detection from whole blood in the integrated microRAAD using 21-day dried amplification reagents. As expected, the red blood cells remained in the MF1 as a spot below the sample inlet while the remaining plasma and buffer solution with virus migrated to the PES for amplification. Following the reaction, we visually observed positive test bands on the LFIAs within 5-10 minutes after valves opened. There was a statistically significant difference between the test band intensity of the 10^5^ virus copies/reaction compared to the negative control (0) (p-value < 0.01) (Figure 4A and C). Previous groups have reported 5 to 10-fold reductions in sensitivity when translating manual assays into automated sample-to-answer assays (Horst et al., 2018; Lafleur et al., 2016; Rodriguez et al., 2015). However, we demonstrate a similar sensitivity using microRAAD for HIV-1 viral detection in blood using dried reagents compared to a standard tube reaction with similar conditions (Figure 4C and Figure 3B). Liu et al. designed a device to detect viral RNA from oral fluid samples in real time down to 12.5 virus copies/reaction, however, viral lysis is required before sample addition and is followed by four more manual steps prior to initiation of the RT-LAMP assay (Liu et al., 2011). Damhorst et al. developed a microfluidic chip for blood cell lysis and modified a smartphone for real-time detection of HIV-1 virus with an LOD of 1.7 × 10^4^ virus copies/reaction (Damhorst et al., 2015). However, the user is required to transfer the lysed blood from the microfluidic chip and freshly prepared RT-LAMP reagents to the reaction chamber for amplification (Damhorst et al., 2015). Even though this platform is 10-fold more sensitive than microRAAD, we believe that the full automation of microRAAD, which reduces sample handling and exposure bloodborne pathogens, makes it an advantageous system for rapid HIV testing at the point of care.

Our initial studies of this integrated sample-to-answer device demonstrate its potential to provide simple, affordable, and early detection of HIV from blood samples at the point of care. The consumable components of microRAAD (membranes, LFIA, adhesive) cost only $2.23 per assay (Table S4) while the reusable components (temperature control circuit and housing) are $70.08 and expected to decrease with increased production (Table S5). The price is comparable with other rapid HIV tests developed for resource-limited settings and will decrease as we scale-up the manufacturing of the device (Rouet and Rouzioux, 2007). Even though microRAAD has many advantages over comparable diagnostic tools, there remain some limitations. The sensitivity of this prototype is 10^5^ virus copies/reaction, equivalent to 4 × 10^6^ virus copies/mL, which falls at the high end of the clinical range, 10^7^ virus copies/mL at peak infection at day 17 (Pilcher et al., 2007). We expect that improvements in primer design and the addition of virus concentration, e.g. size-based capture of virus in a smaller pore-sized membrane, could improve the device’s sensitivity and further enhance clinical relevance. Additionally, microRAAD lacks an internal amplification control to differentiate negative from invalid results (Horst et al., 2018). Incorporating these improvements will enable clinically relevant detection and early diagnosis of HIV.

## Conclusions

We have demonstrated an autonomous and fully integrated sample-to-answer device, microRAAD, for the detection of HIV-1 virus from human whole blood. After sample addition, the LFIA can be visualized within 90 minutes. Moreover, the user is required to perform only four steps to initiate the testing: load sample with rehydrating mixture, add wash buffer, seal the inlets with adhesive, and initiate the temperature control circuit by connecting a power source such as a computer, cellphone, or portable battery. One of the most noteworthy aspects of microRAAD is the complete automation from blood-in to results-out; requiring no sample preparation by the user. Furthermore, we have developed a novel RT-LAMP assay and proved that reagents can be dried and stored at room temperature for three weeks before use in the integrated device. The ability to dry reagents eliminates the need for cold chain storage and increases the usability and portability of the device, especially in resource-limited settings. The sensitivity of this prototype is 10^5^ virus copies/reaction, equivalent to 4 × 10^6^ virus copies/mL, which is below the clinically reported HIV-1 concentration at the peak of infection (Pilcher et al., 2007).

MicroRAAD also has the potential to serve as a platform for detection of other pathogens. By modifying the LAMP primers to target a new gene of interest and adjusting sample preparation depending on the sample matrix, this platform could be used for other viruses (e.g. DENV, CHIKV), and even bacteria (e.g. *Escherichia coli, Vibrio cholerae, Bordetella pertussis*) and parasitic (e.g. *Plasmodium falciparum*) pathogens. This rapid, integrated, and automated device lends itself to use in low-resource areas where clinics and laboratory resources are scarce and gold-standard testing can take up to one week (Wu and Zaman, 2012). Moreover, microRAAD requires only $2.23 worth of consumable components, making it an affordable detection tool. MicroRAAD combines sensitive molecular techniques with elegant capillary fluidics and resilient heating controls into a single, portable platform for rapid pathogen detection at the point of care.

## Acknowledgements

This work was funded by the Grand Challenges Explorations Program (OPP1150806), an initiative of the Bill & Melinda Gates Foundation, the National Science Foundation Graduate Research Fellowship Program (DGE-1333468) (EAP), the National Institute of Allergy and Infectious Diseases (R61AI40474), Purdue University’s Shah Family Global Innovation Lab, and the Purdue Institute for Inflammation, Immunology and Infectious Disease (PI4D).

