## Supplementary Information for "Microfluidic Rapid and Autonomous Analytical Device (microRAAD) to Detect HIV from Whole Blood Samples"

### Electronic Supplementary Information

**Table S1.** Nucleotide sequences of LAMP primers that target the *gag* gene.

| Primer | Sequence (5' – 3') |
|---|---|
| B3 | AGTTCCTGCTATGTCACTTC |
| F3 | TCAGCATTATCAGAAGGAGC |
| BIP | ATGAGGAAGCTGCAGAATGGGCCCTTGGTTCTCTCATCTG |
| FIP | GGTCTCTTTTAACATTTGCATGGCTTTAAACACCATGCTAAACACA |
| LB | AGTGCATGCAGGGCCTATTG |
| LF | TGCTTGATGTCCCCCCAC |
| LB-FITC | /56-FAM/AGTGCATGCAGGGCCTATTG |
| LF-Biotin | /5-Biosg/TGCTTGATGTCCCCCCAC |

**Table S2.** RT-LAMP master mix used for amplification of HIV.

| **Reagent** | **Concentration** |
|---|---|
| Isothermal Buffer II | 1.0X |
| dNTPs | 1.5 mM |
| Betaine | 200 mM |
| F3 Primer | 0.2 µM |
| B3 Primer | 0.2 µM |
| FIP Primer | 1.6 µM |
| BIP Primer | 1.6 µM |
| LB Primer | 0.8 µM |
| LF Primer | 0.8 µM |
| EvaGreen Dye | 0.2X |
| ROX Reference Dye | 1.0X |
| Bst 3.0 Polymerase | 4 U |
| Sucrose | 165 mM |
| Glycerol | 0.28% |
| Triton X-100 | 0.007% |
| Sample | 2-4 µL |
| DEPC $H_2O$ | Fill to 25 µL |

**Table S3.** HIV RT-LAMP mixtures for reagent drying.

| Primer Mixture | Enzyme Mixture | Rehydrating Mixture |
|---|---|---|
| 44.8 mM Sucrose<br>0.007% Triton X-100<br>0.28% Glycerol<br>0.2 µM F3/B3<br>1.6 µM FIP/BIP<br>0.8 µM LF/LB | 120 mM Sucrose<br>1.5 mM dNTPs<br>4 U Bst 3.0 Polymerase | 1X Isothermal Buffer II<br>200 mM Betaine |

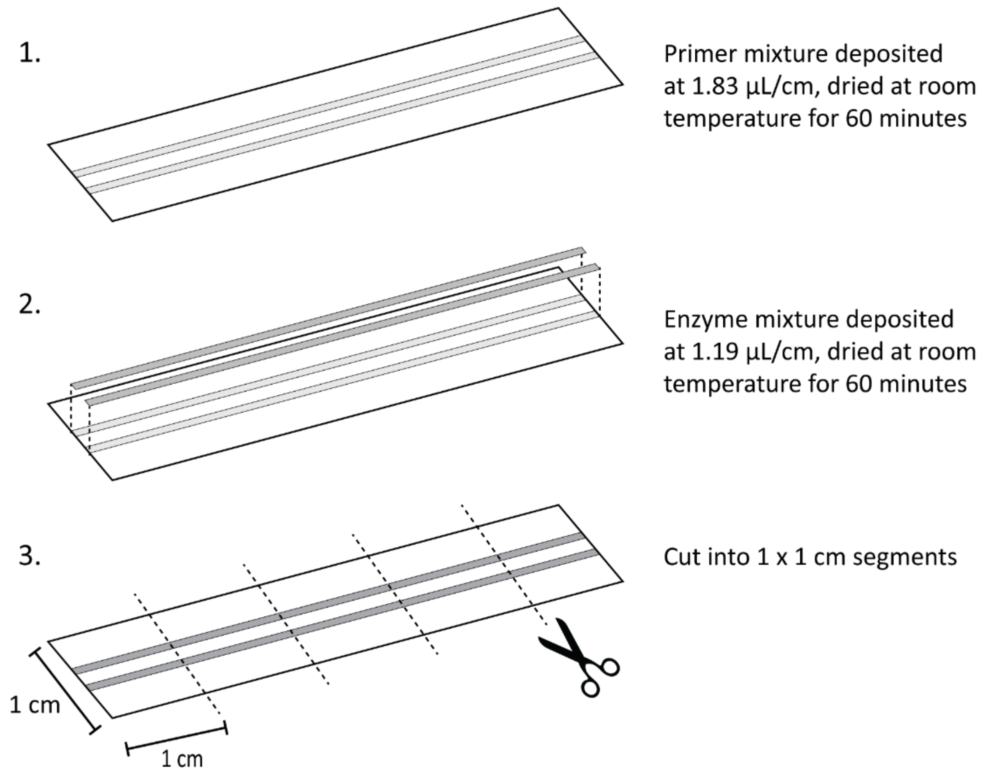


**Figure S1.** HIV RT-LAMP reagent drying setup. The primer mixture is deposited onto PET in parallel lines and let dry at room temperature under continuous air flow for 60 minutes. The enzyme mixture is then deposited directly on top the dried primer mixture and let dry for another 60 minutes at room temperature under continuous air flow. PET with deposited dried reagents is then cut into 1 x 1 cm segments which corresponds to one 25 µL reaction.

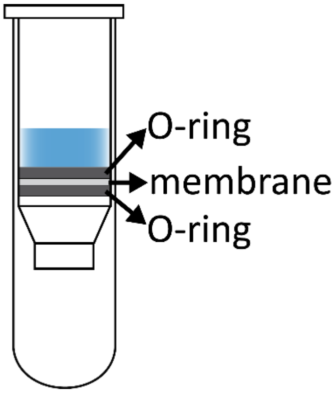


**Figure S2.** Vertical flow filtration setup. Membrane of interest was placed between two O-rings (after removing commercial filter in Qiagen spin column) and placed into spin column before solution was added.

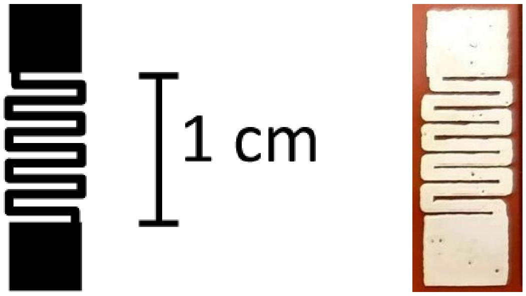


**Figure S3.** Resistive heating elements. Design and image of the resistive heating element after the printing and curing process.

**Table S4.** Components and cost of the consumable components of microRAAD.

| | Component | Manufacturer | Cost/Device |
|---|---|---|---|
| μPAD | Glass fiber | Millipore | $ 0.02 |
| | MF1 blood separator | GE Life Sciences | <$0.01 |
| | 0.22 μm polyethersulfone (PES) | Millipore | $ 0.03 |
| | Wax valve strips | Whatman & Xerox | <$ 0.01 |
| | Cellulose L | Whatman | <$ 0.01 |
| | polyethylene terephthalate (PET) | Apollo | <$ 0.01 |
| | LFIA | USTAR | $ 1.80 |
| | Laminate | Swingline SelfSeal | $ 0.05 |
| | polystyrene gasket | Lohmann Precision Die Cutting | $ 0.01 |
| | Double-sided adhesive | Silhouette | <$ 0.01 |
| | | ***Subtotal*** | ***<$ 1.96*** |
| RT-LAMP Reagents | Isothermal Buffer II | New England Biolabs | $ 0.03 |
| | dNTPs | Agilent Technologies | $ 0.05 |
| | Betaine | Millipore Sigma | $ 0.03 |
| | Primers | Integrated DNA Technologies | $ 0.01 |
| | Bst 3.0 Polymerase | New England Biolabs | $ 0.14 |
| | Sucrose, Glycerol, TritonX-100, green dye, DEPC H2O | Various | $ 0.01 |
| | | ***Subtotal*** | ***$ 0.27*** |
| | | **Consumable Total** | **$ 2.23** |

**Table S5.** Components and cost of the reusable components of microRAAD.

| | Component | Manufacturer | Cost/Device |
|---|---|---|---|
| Resistive heating elements | nanosilver ink | Novacentrix | $ 0.03 |
| | Kapton substrate | | $ 0.45 |
| | | ***Subtotal*** | ***$ 0.48*** |
| Temperature Control Circuit | ATmega328P | Microchip Technology | $ 2.14 |
| | Pogo Pins | Mill-Max Manufacturing Corp. | $ 3.96 |
| | MLX90614 | Melexis Technologies NV | $ 44.34 |
| | AP3429 | Diodes Incorporated | $ 0.42 |
| | Micro USB Female | Amphenol FCI | $ 0.46 |
| | Green LED | Lite-On Inc | $ 0.27 |
| | DAC6311 | Texas Instruments | $ 5.22 |
| | N-MOSFET | Infineon Technologies | $ 1.77 |
| | 2.2µH Inductor | Bourns Inc. | $ 0.24 |
| | 6-pin Female Headers, Right Angle | Sullins Connector Solutions | $ 0.66 |
| | 8MHz Crystal | EPSON | $ 0.83 |
| | Tiny Rectangular Button | C&K Components | $ 0.56 |
| | 22µF capacitor | Samsung Electro-Mechanics America, Inc. | $ 1.38 |
| | 10µF capacitor | Samsung Electro-Mechanics | $ 1.26 |
| | 100n capacitor | Samsung Electro-Mechanics America, Inc. | $ 0.70 |
| | 22p capacitor | Samsung Electro-Mechanics America, Inc. | $ 0.30 |
| | 10k resistor | Stackpole Electronics Inc. | $ 0.60 |
| | 0.1Ohm resistor | Panasonic Electronic Components | $ 1.35 |
| | 1k resistor | Bourns Inc. | $ 0.40 |
| | 300k resistor | Yageo | $ 0.10 |
| | 95.3k resistor | Yageo | $ 0.10 |
| | Printed circuit board (PCB) | | $ 0.50 |
| | | ***Subtotal*** | ***$67.56*** |
| Plastic housing | plastic housing | Stratasys | $ 1.89 |
| | acrylic lid | Shape Products | $ 0.15 |
| | | ***Subtotal*** | ***$ 2.04*** |
| | | **Reusable Component Total** | **$ 70.08** |

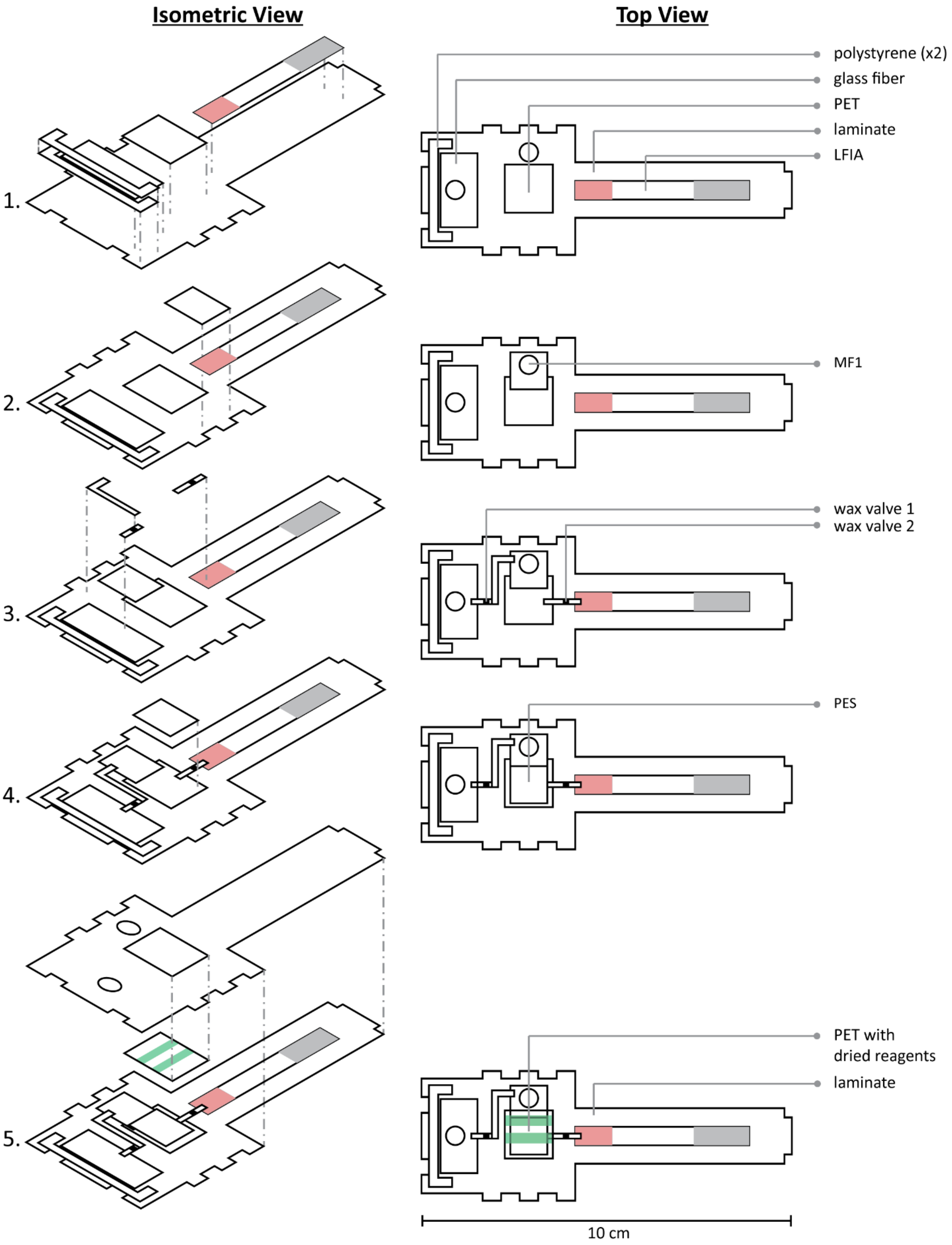


**Figure S4.** Assembly of µPAD. PES was sandwiched with squares of PET to prevent the laminate from inhibiting amplification.

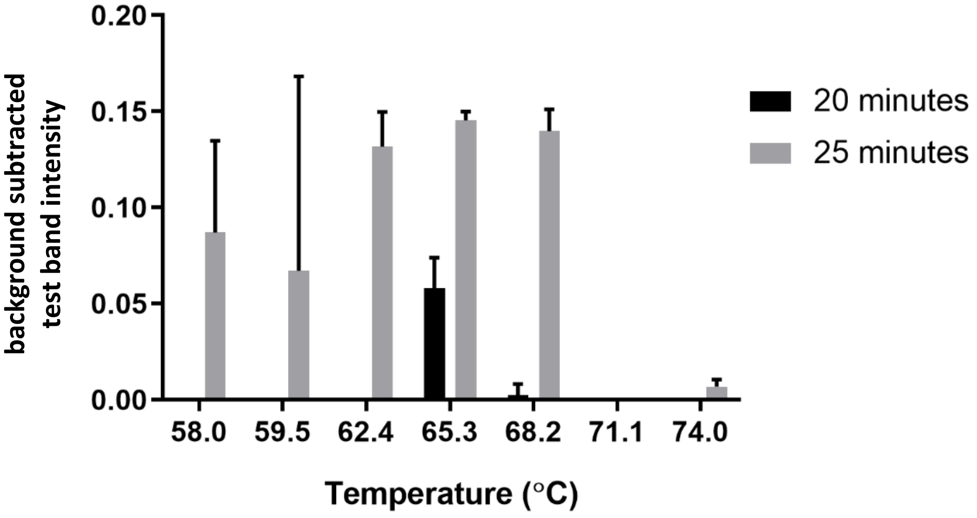


**Figure S5.** RT-LAMP assay efficiency at various temperatures. Amplification products were analyzed by LFIA after 20 and 25 minutes of heating at temperatures ranging from 58 - 74 ºC. When heated for 20 minutes, only amplicons heated at 65 ºC were detectable on LFIAs, indicating HIV RT-LAMP is optimally efficient at 65 ºC. n=2

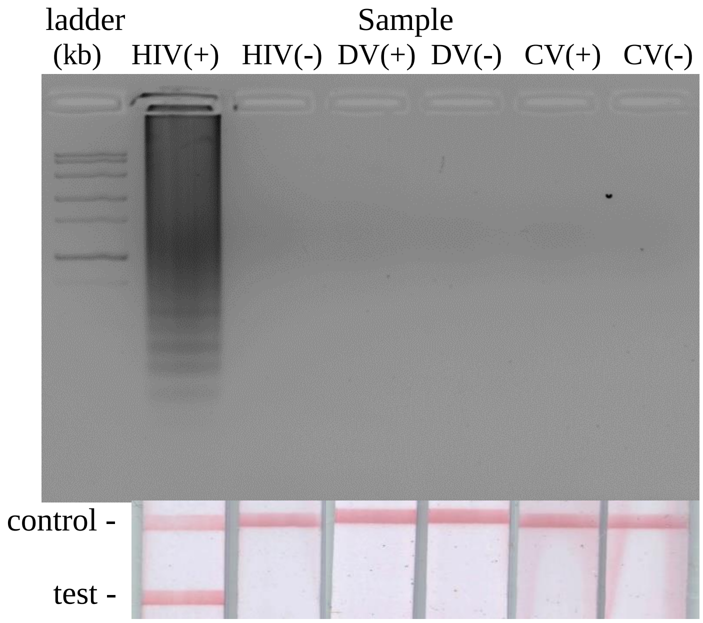


**Figure S6.** Specificity of HIV LAMP primers. Using RNA from Dengue Type 1 Virus (DV) and Chikungunya S-27 Virus (CV) at a concentration of $10^5$ RNA copies/reaction, the HIV RT-LAMP assay was performed for 60 minutes at 65ºC. The only sample that is positive by both gel electrophoresis and LFIA is the HIV(+) sample.

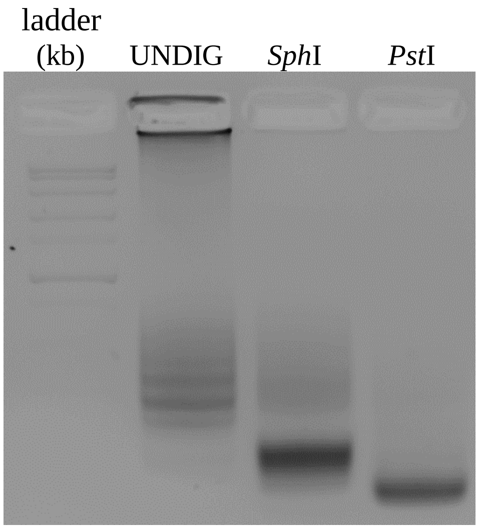


**Figure S7.** Restriction digest of HIV RT-LAMP amplicons. Restriction enzymes *Sph*I and *Pst*I were used to cut the RT-LAMP product, the amplified *gag* gene. Digested products were compared to the undigested product (UNDIG) using gel electrophoresis.

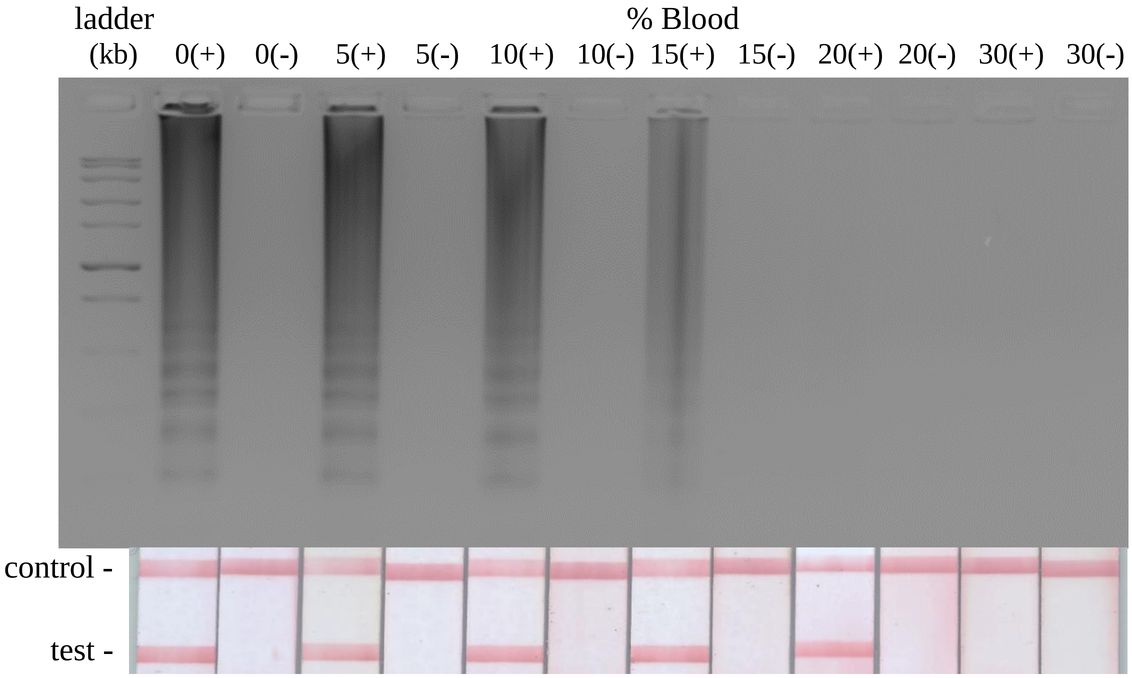


**Figure S8.** HIV RT-LAMP in human whole blood. HIV-1 virus at a concentration of $10^5$ virus copies/reaction was spiked into varying concentrations of blood (0-30%). Gel electrophoresis indicates that the assay tolerance for whole blood is 15% while LFIA demonstrates a tolerance of 20%.

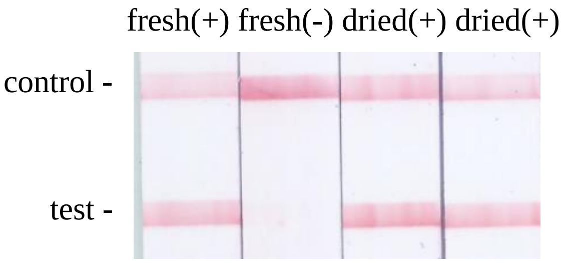


**Figure S9.** Testing 5-month dried RT-LAMP reagents. Dried reagents were rehydrated with $10^6$ virus copies/reaction and rehydrating mixture and amplified for 60 minutes alongside freshly prepared controls.

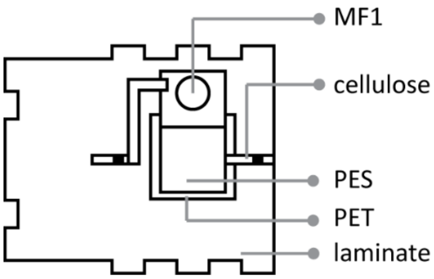


**Figure S10.** Schematic of MF1/PES assembly for studies verifying red blood cell and virus separation.

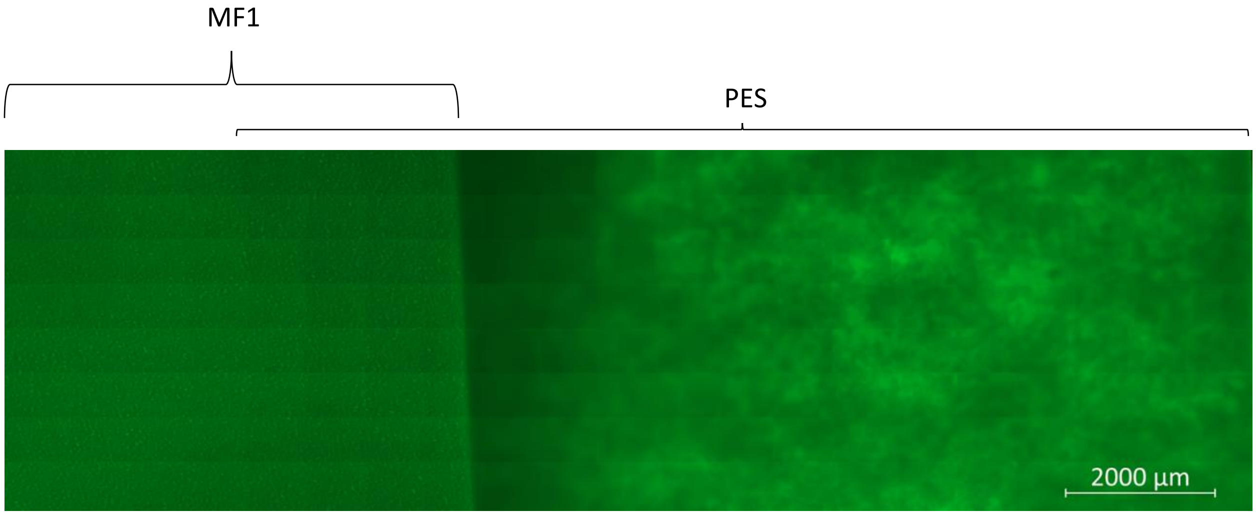


**Figure S11.** Tiled fluorescent images of 100 nm nanoparticles trapped in MF1 and PES of the MF1/PES assembly (Figure S10).

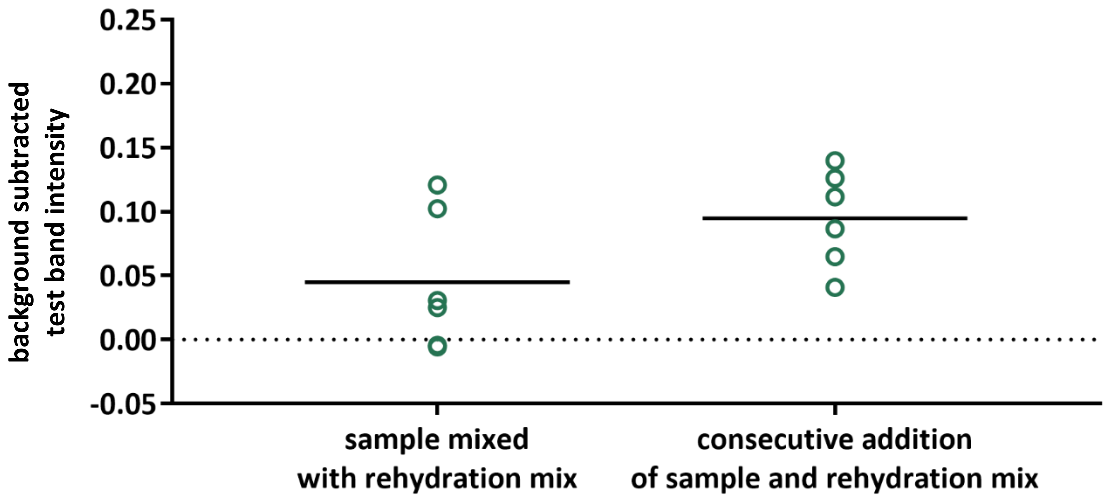


**Figure S12.** LFIA test band intensity of HIV RT-LAMP products after virus and blood cell separation in MF1. $10^5$ copies of HIV virus were diluted in blood and applied to the MF1 of an MF1/PES assembly (Figure S10). Rehydrating mixture was either mixed with the virus in blood and applied simultaneously or applied after the virus in blood (consecutive addition). The PES was removed from the assembly and amplified in RT-LAMP master mix. When the spiked blood was mixed with the rehydrating buffer and applied simultaneously to the MF1 (left), more red blood cells migrated to the PES and inhibited the RT-LAMP assay. n=6.

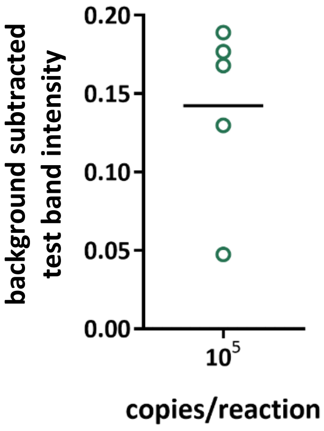


**Figure S13.** Test band intensity of LFIAs when HIV virus was diluted in water and loaded with fresh RT-LAMP reagents into the microRAAD. n=5
